# Salicylic acid contributes to plant defense against a necrotroph: evidence from a transgenic NahG-expressing strain in *Botrytis cinerea*

**DOI:** 10.64898/2026.01.07.698134

**Authors:** Gustavo Hoppe, Antonia Donaire-Guerra, Daniela López-Leiva, Gabriel Pérez-Lara, Francisca Blanco-Herrera, Ariel Herrera-Vásquez, Paulo Canessa

## Abstract

*Botrytis cinerea* is a plant pathogen that causes significant agricultural losses worldwide. Although this necrotroph disrupts extensive plant hormonal networks, the role of salicylic acid (SA) in plant defense against *B. cinerea* remains controversial across plant species. To investigate its role from a pathogen perspective, *B. cinerea* mutants constitutively expressing the *Pseudomonas putida* salicylate hydroxylase NahG, an enzyme that catalyzes salicylic acid degradation, were generated.

The NahG-expressing *B. cinerea* mutants exhibited enhanced *in vitro* growth on SA-supplemented media, indicating that SA catabolism confers an advantage. *In planta*, these mutants displayed increased virulence in *Arabidopsis thaliana* and *Phaseolus vulgaris*. Notably, the increase in lesion formation was strictly dependent on host SA biosynthesis, as no differences were observed when infecting the SA-deficient Arabidopsis *sid2-2* mutant. This result provides evidence that SA degradation increases the virulence of *B. cinerea* in the interaction with *A. thaliana.* Genome inspection revealed that the fungus encodes four salicylate hydroxylase–like genes. Analysis of publicly available transcriptomic data from virulence assays across multiple plant hosts revealed that all these genes are expressed during the plant-pathogen interaction, with distinct expression patterns across infection stages and hosts. Together, these observations suggest that *B. cinerea* may have endogenous mechanisms for SA degradation during host colonization, thereby conferring the capacity to control its accumulation during the infection process.

**HIGHLIGHTS:**

- *Botrytis cinerea* expressing the salicylate hydroxylase (SH) ppNahG shows enhanced virulence in Arabidopsis and bean plants.
- Enhanced virulence of NahG-expressing strains depends on host salicylic acid biosynthesis.
- The *Botrytis cinerea* genome encodes four SH–like genes.
- SH-like genes display distinct expression patterns during infection across different plant hosts.

**VISUAL ABSTRACT:** 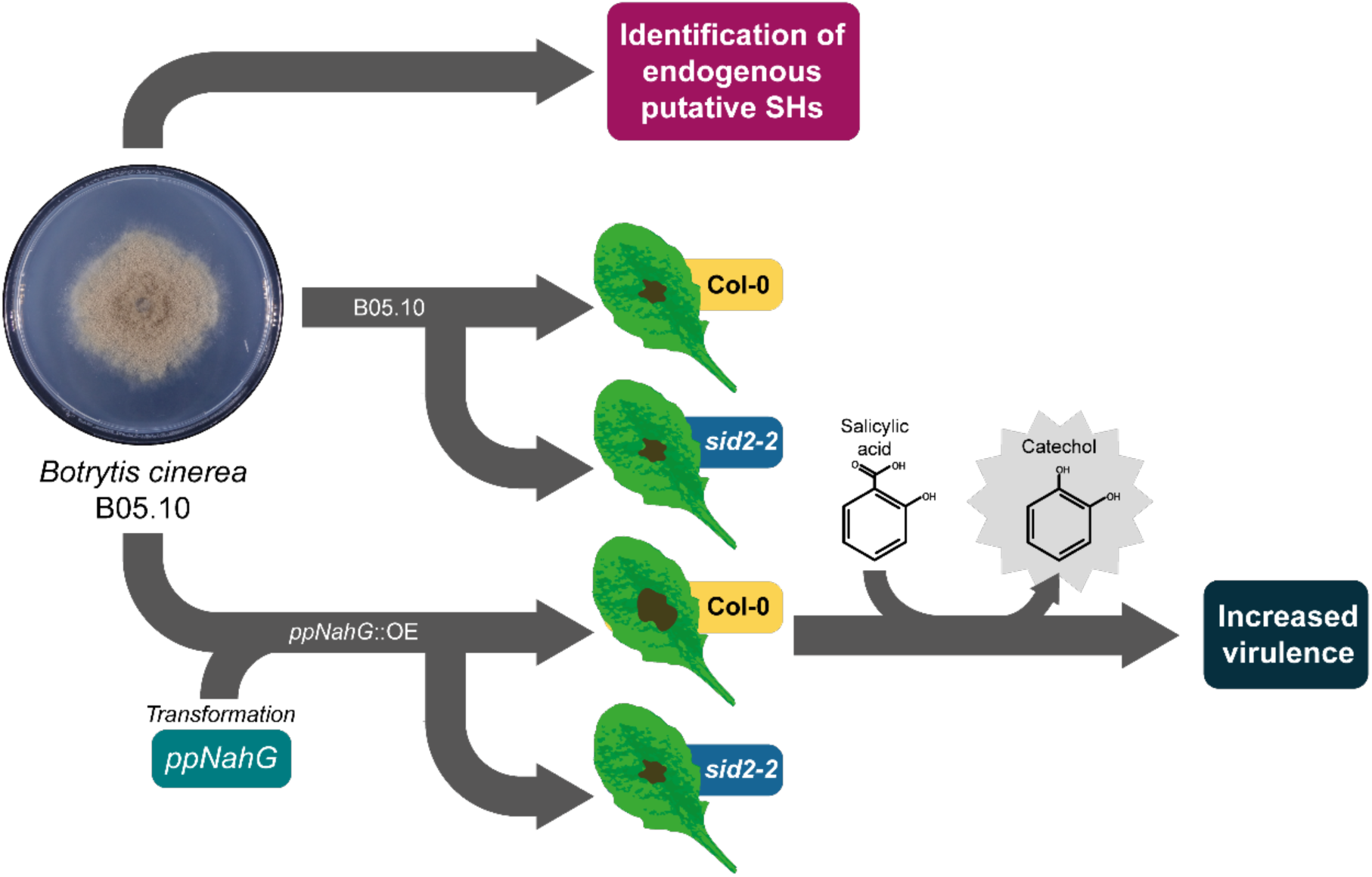

## INTRODUCTION

*Botrytis cinerea* is one of the most notorious necrotrophic fungal pathogens worldwide, causing gray mold disease in over 1,400 plant species, including economically valuable crops such as tomato, grapevine, and strawberry, among others (Elad et al., 2016; Veloso & Van Kan, 2018). It is one of the most economically damaging fungal phytopathogens (Dean et al., 2012). As a necrotroph, *B. cinerea* colonizes plant tissues by inducing host cell death, thereby allowing it to feed on dead plant material (Bi et al., 2023). The infection process of this fungus is complex and involves the secretion of diverse virulence factors, including cell wall–degrading enzymes and phytotoxins. These, among other factors, enable rapid disease progression, making this fungus difficult to control in agricultural settings (Choquer et al., 2007; Shlezinger et al., 2011; Van Kan, 2006).

To establish a successful infection, *B. cinerea* not only confronts but also reprograms host immune responses. A key target in this manipulation is the balance between two defense-related phytohormones known as salicylic and jasmonic acid (SA and JA, respectively) (El Oirdi et al., 2011; Thomma et al., 1998; Wei et al., 2024), both of which are central regulators of plant immunity (Glazebrook, 2005; Hou & Tsuda, 2022). JA, along with ethylene (ET), are considered the canonical signals associated with necrotrophic resistance (Macioszek et al., 2023). The JA-dependent defenses induce the expression of genes encoding peptides with antimicrobial properties, such as Plant Defensin 1.2 (PDF1.2), secondary-metabolite producing enzymes, and proteins that reinforce the plant cell wall (Penninckx et al., 1998). Moreover, exogenous application of JA or methyl jasmonate has been shown to provide a protective effect against *B. cinerea* through peroxide and anthocyanin production (Jiang et al., 2015; Ren et al., 2020; Valenzuela-Riffo et al., 2020; Yang et al., 2025). In this regard, various studies have demonstrated that plants with defects in JA biosynthesis or signaling (e.g., *dde2* and *coi1*, respectively) are more susceptible to *B. cinerea*, highlighting the critical role of JA in defense (AbuQamar et al., 2006; Thomma et al., 1998).

Notably, *B. cinerea* secretes an exopolysaccharide that can trigger SA accumulation in the host. In turn, this activity suppresses JA signaling, a textbook example of a pathogen strategy to manipulate plant defense mechanisms through phytohormone cross-talk (El Oirdi et al., 2011). The pathogen also produces the toxin known as botrydial, which triggers cell death mimicking a hypersensitive response associated with SA, thereby enhancing tissue maceration through inappropriate activation of a biotroph-targeted pathway (Colmenares & Aleu, 2002; Rossi et al., 2011). These observations support the prevailing view that syndicate JA as a necrotroph-protective phytohormone, whereas SA is often detrimental (Glazebrook, 2005; Mengiste, 2012). Yet evidence challenges this binary framework, as several studies demonstrate that SA can contribute to basal resistance against *B. cinerea*. For instance, exogenous SA application or functional analogs such as Benzothiadiazole (BTH) have been shown to reduce lesion size and enhance the expression of pathogenesis-related (PR) genes, such as *PR1,* in both *A. thaliana* and *Solanum lycopersicum* (Audenaert et al., 2002; Ferrari et al., 2003; García-Pastor et al., 2020; Zimmerli et al., 2001). Conversely, *nahG* transgenic plant lines expressing a bacterial salicylate hydroxylase, which depletes SA, consistently exhibit increased susceptibility (Audenaert et al., 2002; Ferrari et al., 2003; Govrin & Levine, 2002). These findings suggest that SA may operate alongside or upstream of JA to restrict necrotrophic growth.

The apparently contradictory roles attributed to SA illustrate the complex nature of hormone-driven plant immunity. The accumulation of SA, whether from exogenous application or biosynthesis, the timing and spatial context of its effects, and tissue- or species-specificity, all affect infection outcome. Besides, all these variables operate within a complex hormonal network that collectively shapes the end product of the infection (Tsuda & Katagiri, 2010). Indeed, it has been demonstrated that viral infections can increase both SA levels and susceptibility to *B. cinerea* (Gupta et al., 2023), while SA-producing rhizobacteria are able to induce systemic resistance in wild-type plants but not in *nahG* transgenic lines (De Meyer & Höfte, 1997). As these outcomes depend on each context, there is a need for studies that go beyond the host’s perspective.

Through the use of advanced genetic tools in plants, our understanding of SA responses and signaling has advanced dramatically. Mutants deficient in the biosynthesis or signaling of this phytohormone, such as *ics1* (*sid2*), *npr1*, and *eds1*, display augmented susceptibility to biotrophic pathogens, although their response to *B. cinerea* is variable (Ferrari et al., 2003; Tsuda et al., 2008). Indeed, increased resistance to *B. cinerea* is reported in *npr1-1* mutant plants, in which SA signaling is compromised. This phenotype can be attributed to a possible reduced suppression of JA signaling. Conversely, *sid2* mutants exhibit enhanced susceptibility, implicating SA in basal defense (Dewdney et al., 2000; Spoel et al., 2003). The use of *nahG* lines has further revealed that complete depletion of SA affects not only *PR* gene expression but also associated responses such as the oxidative burst and cell wall reinforcement (Van Wees et al., 2000).

Beyond signaling, SA has antifungal activity. *In vitro* studies have shown that SA can alter mitochondrial function, thereby inhibiting fungal growth. SA can disrupt membrane integrity and metal homeostasis, promoting the production of reactive oxygen species (ROS) (Qi et al., 2012). Similar fungistatic effects have been documented for other necrotrophs, including *Fusarium graminearum*, *Aspergillus flavus*, and *Harpophora maydis* (Degani et al., 2015; Panahirad et al., 2014; Rocheleau et al., 2019), suggesting a broad antimicrobial function.

To explore whether *B. cinerea* also benefits from depleting SA during infection, we engineered fungal transgenic strains expressing the bacterial salicylate hydroxylase *nahG* from *Pseudomonas putida* (*ppNahG*) (You et al., 1990). Rather than altering host SA metabolism or signaling, this reverse-genetics approach provides a pathogen-centered perspective and a direct means to test the effect of local SA removal on infection dynamics. Accordingly, we examined the *B. cinerea* lesion-forming ability in *A. thaliana* and *Phaseolus vulgaris*, two complementary models, to determine whether SA degradation confers an advantage. Our findings provide new insights into the multifaceted role of SA in plant–necrotroph interactions and reveal a potentially widespread fungal strategy based on hormone degradation.

## RESULTS

To investigate the role of SA during plant infection by a necrotroph, we engineered a *B. cinerea* mutant expressing the *ppNahG* gene from *P. putida*, thereby providing the fungus with a constitutive ability to degrade host-derived SA during infection. For this purpose, a genetic construct was designed and assembled to target integration into the *bcku70 locus* of *B. cinerea* (Choquer et al., 2008) via homologous recombination (Hamada et al., 1994). The construction included the *ppNahG* coding sequence (NCBI Gene ID: X83926.1) under the control of the actin A (*bcactA*; Bcin16g02020) promoter and the tubulin A (*bctubA*; Bcin01g08040) terminator (Schumacher, 2012), along with the hygromycin resistance cassette *hph* for selection **(Figure S1A)**. Two independent mutants, designated *ppNahG*::OE-1 and *ppNahG*::OE-2, were obtained. The correct integration at the mentioned *locus* was confirmed by PCR **(Figure S1B)**.

To assess the effect of *ppNahG* on fungal development, an *in vitro* growth assay was performed employing different carbon sources. Fungal growth was evaluated on the minimal Gamborg B5 medium supplemented with 2% sucrose, 1 mM SA, or 1 mM catechol (the product of SA catabolism by the *ppNahG* enzyme). No significant differences in fungal growth, measured as colony area, were observed between the B05.10 strain and the *ppNahG*::OE mutants when cultured on medium supplemented with 2% sucrose **(Figure 1A and B)**. When SA was provided **(Figure 1A and C)** as the sole carbon source, the mutants expressing the *ppNahG* enzyme (*ppNahG*::OE-1 and *ppNahG*::OE-2) exhibited greater growth area after 7 days when compared to the B05.10 strain, suggesting increased capacity to degrade SA. No significant differences were observed when catechol was provided as the sole carbon source **(Figure 1A and D)**. These results indicate that *ppNahG* expression increases growth capabilities on SA-containing medium, thereby reducing its antifungal activity, and/or from the capacity to metabolize SA as a carbon source.

**Figure 1.**
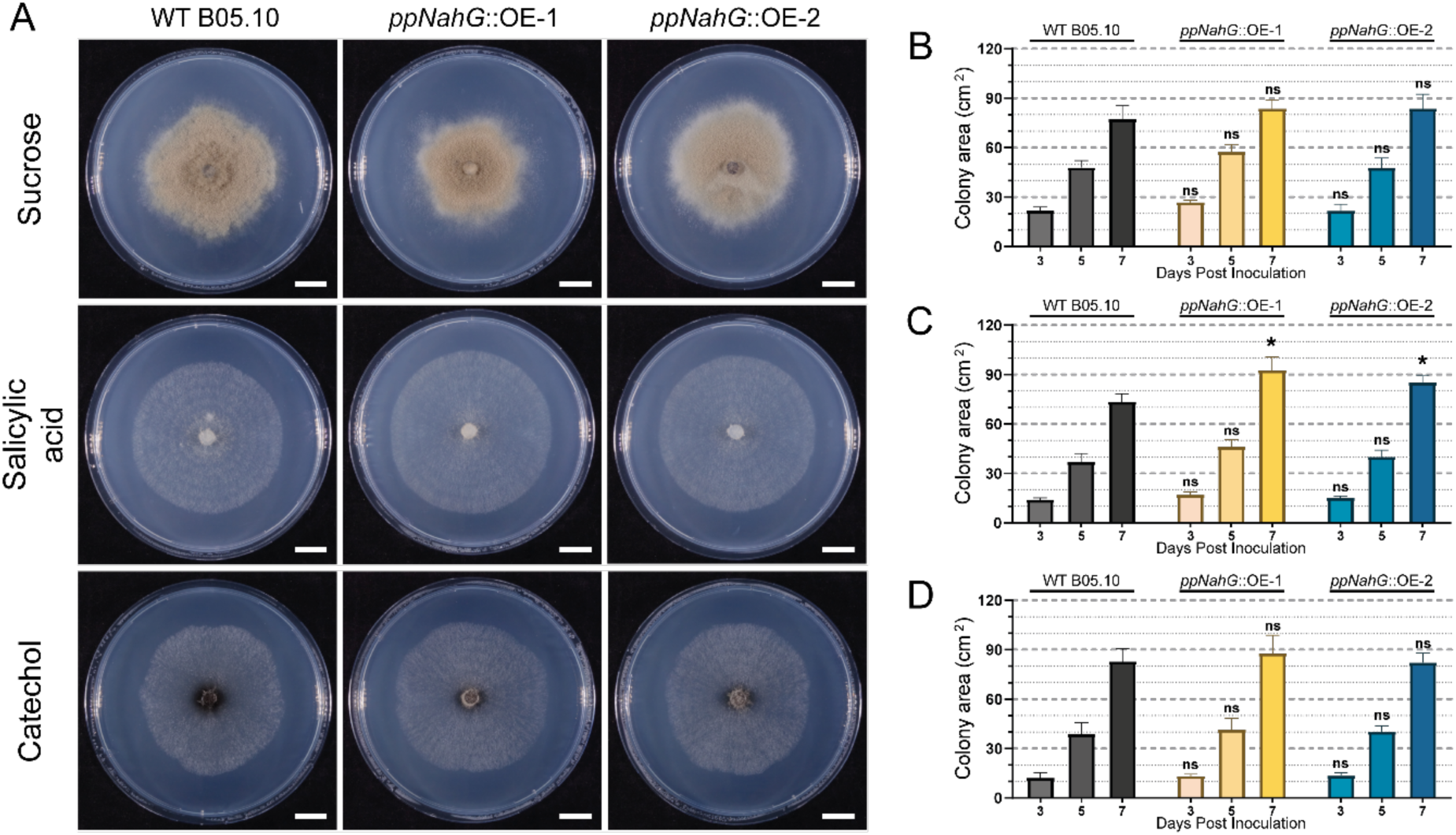
Heterologous expression of a salicylate hydroxylase encoding gene in *Botrytis cinerea* B05.10 enhances the ability to grow in a medium supplemented with salicylic acid. Growth profiles of the wild-type B05.10 strain and two independent *ppNahG::OE* transformants were evaluated 7 days post inoculation (dpi) on solid Gamborg B5 medium supplemented with 2% sucrose, 1 mM salicylic acid, or 1 mM catechol (**A**; scale bar = 2 cm). Colony diameters were quantified at 3, 5, and 7 dpi to determine the effects of sucrose **(B)**, salicylic acid **(C)**, and catechol **(D)** on fungal growth. Error bars indicate the standard deviation from three independent experiments. Statistical significance compared to the control B05.10 strain was determined at each time by T-test (ns, no significant differences, *: *p* < 0.05).

To evaluate the impact of *ppNahG* expression on virulence, *P. vulgaris* and *A. thaliana* plants were infected with the *ppNahG*::*OE* mutants. *P. vulgaris* was selected due to its large leaves and rapid growth, which allowed an initial and visual assessment of fungal virulence in a highly susceptible host. Leaves were inoculated with a conidial suspension of the different strains. The *ppNahG-*expressing mutants caused significantly larger lesions than the B05.10 strain **(Figure 2A)**. Virulence assays were also performed on four-week-old *A. thaliana* Col-0 plants. Plants were inoculated with the B05.10 strain and the *ppNahG::OE* mutants as described (Canessa et al., 2013; Schumacher, 2012). Both mutants induced a statistically significant increase in lesion size, reaching approximately 1.8-fold that of lesions caused by the B05.10 strain **(Figure 2B)**.

**Figure 2.**
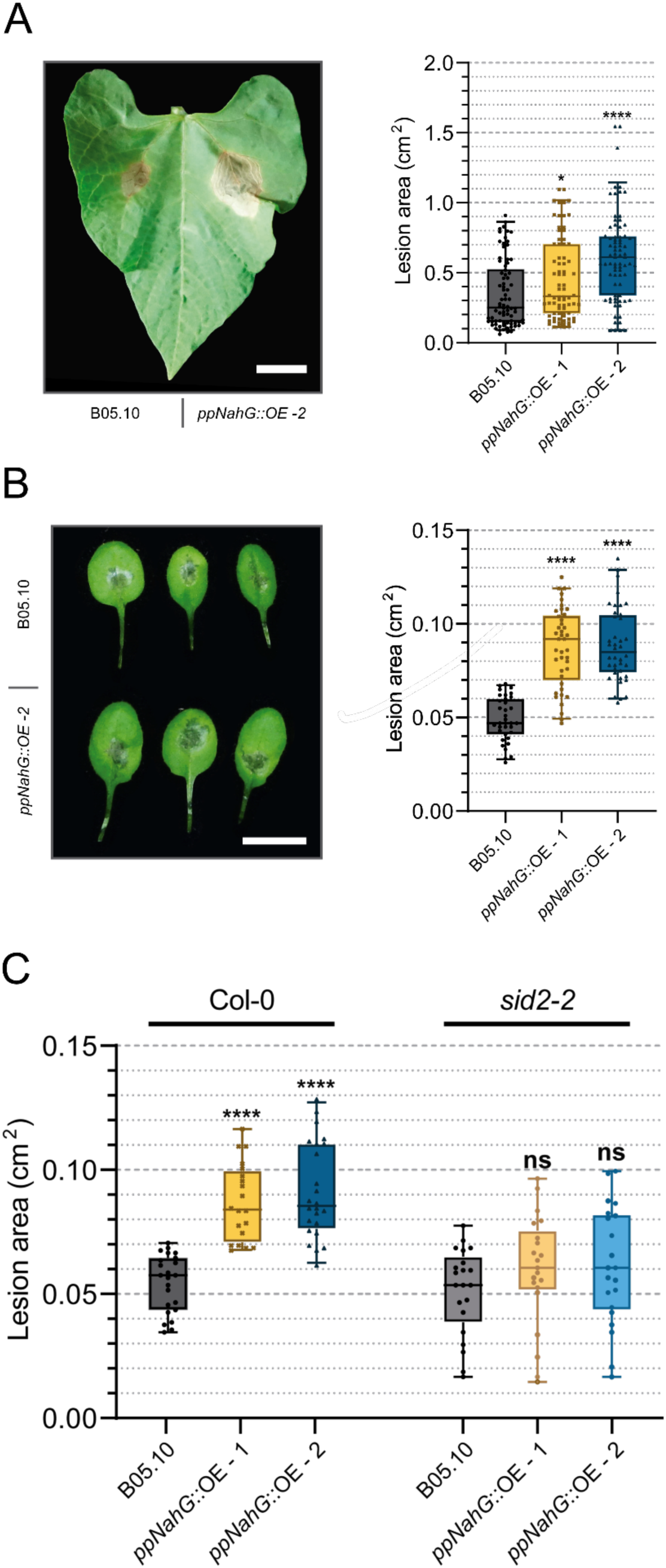
The expression of the *ppNahG* gene increases the virulence of *Botrytis cinerea* in two different plant hosts. **(A)** Virulence assays on *P. vulgaris*. Primary leaves of 1-week-old plants were inoculated with each *ppNahG::OE* mutant and the B05.10 strain. Lesions were photographed 96 hours post inoculation (hpi). A representative leaf is shown (scale bar = 1 cm). The lesion-size quantification is indicated at the right. **(B)** Virulence assays were conducted on *A. thaliana*. Leaves of 4-week-old plants were inoculated as in the Methods. Photographs were acquired at 72 hpi to quantify lesion size. For each fungal genotype, three representative leaves are shown (scale bar = 1 cm). **(C)** Virulence assays conducted on *A. thaliana* Col-0 and *sid2-2* mutant plants. Lesion quantification was performed 72 hpi. In all plots, error bars represent the 95% confidence interval. Statistical significance compared to the control B05.10 strain was determined by ANOVA and Holm-Šídák’s multiple comparisons test (ns, not significant; *: *p <* 0.05; ****: *p* < 0.0001).

To determine whether the virulence differences observed between the B05.10 strain and the *ppNahG::OE* mutants depended on the host’s ability to accumulate SA, *sid2-2 A. thaliana* mutant plants were used. This *sid2-2* allelic mutant lacks a key enzyme in the SA biosynthesis pathway, isochorismate synthase 1 (ICS1) (Wildermuth et al., 2001). The B05.10 strain exhibited similar virulence in both Col-0 and *sid2-2* Arabidopsis, while the *ppNahG*::OE mutants, despite inducing significantly larger lesions on Col-0 plants, showed no difference in virulence against the B05.10 strain when infecting *sid2-2* plants **(Figure 2C)**.

The differential virulence of *ppNahG*::*OE* mutants observed between Col-0 and *sid2-2* plants indicates that IC-mediated SA accumulation impacts lesion development, suggesting that its degradation by ppNahG activity could modulate lesion expansion. For this reason, we analyzed the expression of critical defense responsive genes in Arabidopsis.

To determine whether *ppNahG* expression alters the SA-responsive gene activation, we first analyzed the transcriptional activity of the *PR1* promoter using *A. thaliana* plants expressing YFP under its control (pPR1::YFP). Fluorescence microscopy revealed that *PR1* activation can be detected in cells close to the infection site, with no detectable signal in distal tissue on the same infected leaf **(Figure 3A)**. Importantly, YFP fluorescence was observed in plants infected with the B05.10 strain as well as those infected with the *ppNahG*::OE-2 mutant **(Figure 3A)**. This indicates that *ppNahG* expression does not alter the spatial or qualitative pattern of *PR1* promoter activation during early infection by *B. cinerea* **(Figure 3A** and **Figure S2)**.

**Figure 3.**
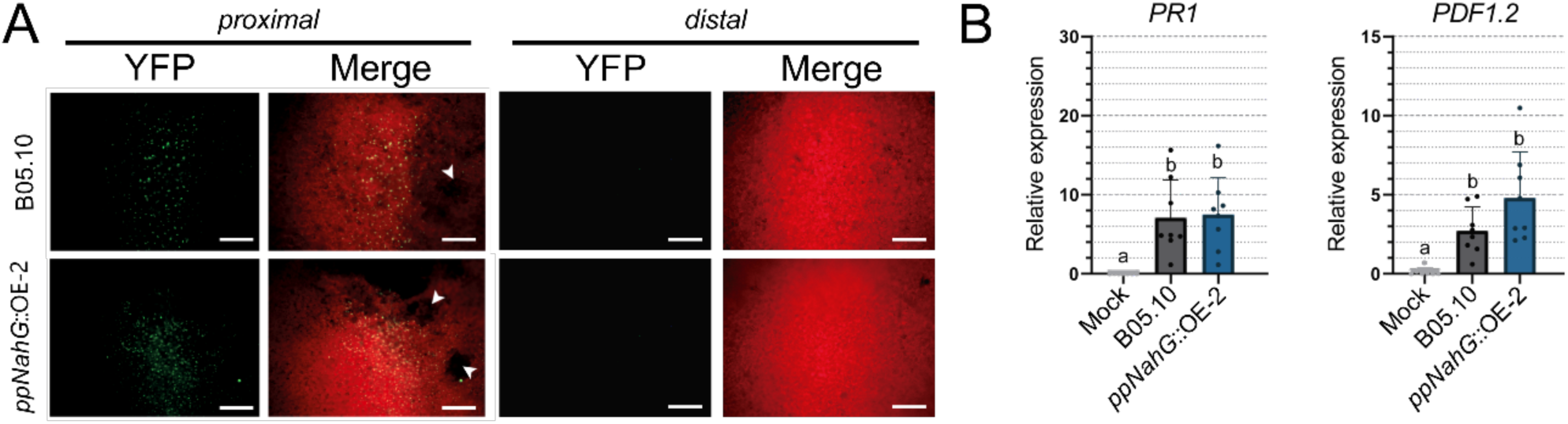
The constitutive expression of *ppNahG* in *Botrytis cinerea* does not affect *PR1* and *PDF1.2* accumulation during the infection of *A. thaliana*. (**A**) Epifluorescence microscopy of *A. thaliana* pPR1::YFP leaves at 24 hours post infection (hpi) with *B. cinerea*. YFP fluorescence is shown in green, and chlorophyll autofluorescence in red. Representative images of proximal and distal regions relative to the lesion site (white arrows) are shown. Scalebar = 50µm. **(B)** Relative expression of *PR1* and *PDF1.2* was measured by RT–qPCR at 24 hpi. *A. thaliana* Col-0 and *sid2-2* plants were inoculated with the B05.10 strain or the *ppNahG* mutant (*ppNahG::OE-2*) of *B. cinerea*. Different letters indicate statistically significant differences by ANOVA and Holm-Šídák’s multiple comparisons test (*p* < 0.05).

To complement these observations, we quantified the accumulation of *PR1* and *PDF1.2* transcripts by RT-qPCR in the proximal infected area of *A. thaliana* leaves. Both *PR1* and *PDF1.2* transcripts increased significantly at 24 hours post inoculation (hpi) compared with mock-inoculated plants, as expected **(Figure 3B)**. However, no differential expression was detected between infections caused by the B05.10 strain and the *ppNahG*::OE-2 mutant. Together, the *PR1* reporter assays and transcript quantification indicate that *ppNahG* expression does not modify SA-responsive gene activation, either at the level of promoter activity or mRNA accumulation, and therefore, is unlikely to underlie the enhanced virulence observed in *ppNahG*::OE mutants.

Although our analyses did not reveal changes in SA-responsive gene expression, the *ppNahG*::*OE* mutants consistently exhibited enhanced virulence. This suggests that pathogen-mediated SA degradation can contribute to disease development. Notably, several microorganisms (Li et al., 2017; Lowe-Power et al., 2016), including fungal pathogens (He et al., 2023; Lubbers et al., 2021; Xu et al., 2025), possess enzymes capable of degrading SA. Moreover, as shown in **Figure 1**, *B. cinerea* B05.10 can grow on SA as the sole carbon source, indicating that the fungus must possess metabolic mechanisms to catabolize SA.

Based on these precedents, we examined whether *B. cinerea* encodes endogenous salicylate hydroxylase–like enzymes that could participate in SA metabolism and potentially contribute to its virulence. Accordingly, we identified four genes encoding proteins with over 25% identity to ppNahG: Bcin06g05670, Bcin05g08190, Bcin10g05120, and Bcin02g00013, with identities of 42.2, 33.6, 31.5, and 26.5 %, respectively. These hypothetical proteins show even higher identity to SsShy1, a salicylate hydroxylase (SH) from *Sclerotinia sclerotiorum* (83.2%, 40.7%, 35.6%, 28.4% identity, respectively) (He et al., 2023). In particular, Bcin06g05670 and Bcin05g08190 are closely related to other fungal SHs with experimentally validated enzymatic activity, strengthening the possibility of SH activity **(Figure 4A)**.

**Figure 4.**
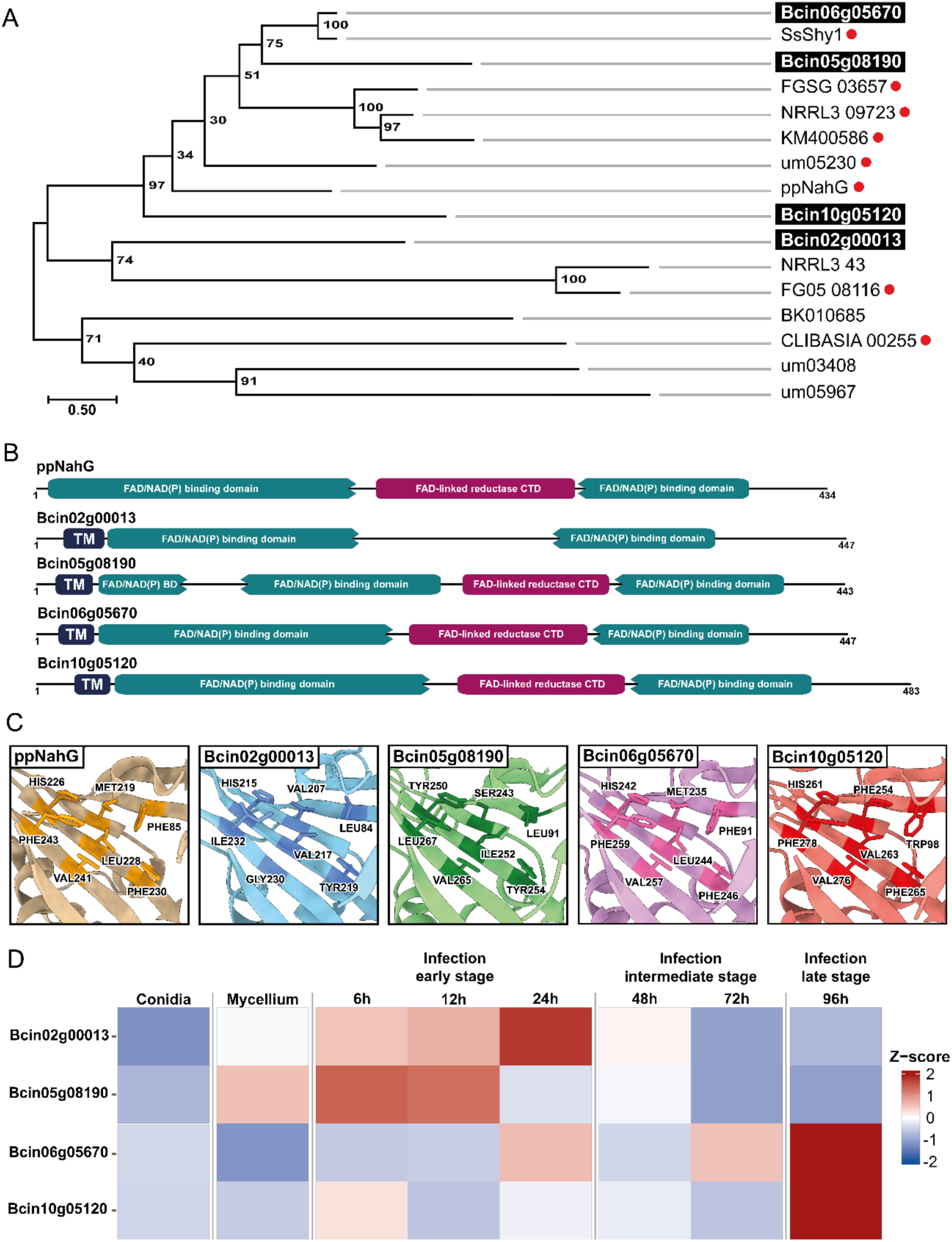
Identification of four putative salicylate hydroxylases in the *Botrytis cinerea* genome. **(A)** Phylogenetic reconstruction of candidate salicylate hydroxylase (SH) proteins encoded in the *B. cinerea* genome. Previously characterized SH proteins from other fungal species, as well as ppNahG, were included for comparison. Enzymes with experimentally verified SH activity are marked with red circles. **(B)** Functional domain predictions for ppNahG and the *B. cinerea* SH candidates. TM denotes transmembrane domains. **(C)** Comparison of the experimentally determined crystal structure of ppNahG (PDB: 6BZ5) with the predicted structures of the four putative *B. cinerea* SH proteins. Active sites and predicted catalytic amino acids are indicated. Predicted structures were obtained from the AlphaFold database (Jumper et al., 2021). **(D)** Heatmap illustrating the relative expression levels of putative SH-encoding genes in *B. cinerea* during *A. thaliana* infection at different timepoints (h). Expression from conidia and mycelium is included. Data obtained from the Sequence Read Archive (SRA) accession number PRJNA1158549 (Wei et al., 2024).

Prediction of functional domains using InterProScan confirms that all but one of the encoded proteins harbor the characteristic domains of SH enzymes, including a FAD-binding domain and a FAD-linked reductase C-terminal domain. These predicted proteins also contain a putative transmembrane domain that is absent in the ppNahG protein (**Figure 4B)**. The active site of the crystallized ppNahG (Costa et al., 2019) was compared with the predicted structure of each candidate as determined by AlphaFold (Jumper et al., 2021). All the amino acid residues previously characterized in ppNahG for the binding of SA (F85, M219, H226, L228, F230, V241, and F243) (Costa et al., 2019) are conserved in analogous positions in the predicted product of Bcin06g05670 **(Figure 4C)**. The product of Bcin10g05120 shared residues equivalent to H226, F243, V241, and F230 of ppNahG, whereas Bcin02g00013 and Bcin05g08190 retained only the residues corresponding to H226 and V241, respectively **(Figure 4C)**. Examining the expression patterns of the identified genes through publicly available transcriptomics data can provide insights into their potential roles during infection **(Figure 4D)**. During the early stages of *A. thaliana* infection by B05.10 (Wei et al., 2024), Bcin02g00013 and Bcin05g08190 exhibit higher expression levels compared to Bcin06g05670 and Bcin10g05120. In contrast, these two genes show their highest expression levels after 96 hpi. This implies that these genes could play a role in the *B. cinerea* infection process, an idea reinforced by their reduced expression in conidia or mycelium, outside of the host colonization context **(Figure 4D)**. Examination of additional transcriptomics data from virulence assays conducted in *S. lycopersicum* and *Lactuca sativa* indicates similar expression patterns **(Figure S3)** (Pink et al., 2023; Srivastava et al., 2020), with increased expression of Bcin02g00013 during early infection and upregulation of Bcin10g05120 and Bcin05g08190 at later time points. In contrast, Bcin06g05670 displays a less consistent expression pattern across the data set, suggesting a regulation influenced by other factors such as host context or other particular experimental conditions. The obtained data further support not only the importance of SA during *B. cinerea* infection, but also the possibility that it has the intrinsic capacity to modulate its accumulation.

## DISCUSSION

Research has explored the role of SA in various plant physiological processes, particularly in plant defense. Still, its function during the infection by a necrotrophic phytopathogen remains comparatively less understood. In the case of *B. cinerea* infections, its role is even harder to pin down. In *A. thaliana*, exogenous SA application protects against *B. cinerea* (Ferrari et al., 2003), whereas the opposite effect is seen during infection of tomato leaves (El Oirdi et al., 2011). This is further complicated by evidence showing that SA application can reduce the damage caused by *B. cinerea* in tomato fruits (García-Pastor et al., 2020), suggesting that the role of SA during infection may depend not only on the host but also on the tissue being infected. This might explain the lack of consensus in the SA-mediated responses against this pathogen.

As in any two-sided story, *B. cinerea* can also affect the SA response in plants. Secreted virulence factors known as cell death inducing proteins (CDIPs), such as BcSPL1 and BcHIP1, not only are capable of generating damage on leaves but also induce SA accumulation when these proteins are heterologously produced and infiltrated into the apoplast (Frías et al., 2013; Jeblick et al., 2023). These observations suggest that SA could be relevant for lesion development. Moreover, the xylanase BcXYL1 induces the expression of the *PR-1a* defense gene in *Nicotiana benthamiana* even when its enzymatic activity is impaired, indicating that this protein is involved in activating the plant immune response independently of its catalytic function (Yang et al., 2018). However, despite these associations between CDIPs and SA-related responses, the functional significance of SA to either lesion formation or host defense during *B. cinerea* infection remains elusive and most likely, context-dependent.

Additional *B. cinerea* virulence factors also appear to act in an SA-dependent manner. For instance, the sesquiterpene botrydial induces lesion formation in *A. thaliana* and tobacco leaves, and its effect is reduced in NahG tobacco plants (Rossi et al., 2011). Furthermore, the production of an exopolysaccharide by the *B. cinerea* strain B191 induces SA accumulation, thereby enabling increased virulence in tomato plants (El Oirdi et al., 2011). Together, these reports indicate that SA-related pathways are tightly intertwined with *B. cinerea*–induced cell death, even though the precise causal role of SA itself remains unclear.

Our results demonstrate that a gain-of-function through the expression of *ppNahG* in *B. cinerea* leads to higher virulence in *P. vulgaris* and *A. thaliana* leaves. Moreover, as demonstrated in *Arabidopsis*, this effect is dependent on the plant’s ability to accumulate SA, as *sid2-2* mutants show no differential susceptibility to *ppNahG::OE B. cinerea* mutants. This observation is noteworthy, as it is consistent with a previous report in which transgenic plants expressing NahG were more susceptible to *B. cinerea*, whereas *sid2-2* plants were not (Ferrari et al., 2003). Importantly, most previous studies have manipulated SA levels at the plant level, either by impairing SA biosynthesis or by exogenous application of SA at high concentrations (El Oirdi et al., 2011; Ferrari et al., 2003; García-Pastor et al., 2020; He et al., 2023), a strategy that affects the systemic response of the plant and does not allow for the evaluation of more localized processes such as initial infection development. In contrast, our approach modifies the pathogen itself to confer SA-degrading capacity, allowing SA turnover to occur at the host– pathogen interface and providing a complementary, pathogen-centered perspective on how SA catabolism can influence disease development by a necrotrophic pathogen.

Collectively, our results indicate that the ability to degrade SA — rather than the mere presence or absence of its signaling — is a determinant of virulence. Indeed, this interpretation is informed by the transcriptional PR1 reporter and RT-qPCR analysis, which revealed no evidence for a major shift toward canonical SA-responsive gene activation. Naturally, PR1 is an indirect single SA marker gene, and therefore, we cannot exclude subtler or transient differences in SA signaling.

In addition to its potential effects on host signaling, SA catabolism may confer a direct metabolic advantage to the pathogen. Consistent with this idea, ppNahG-expressing strains display enhanced growth in the presence of SA, suggesting that degradation of this compound may alleviate its inhibitory effect and/or allow its use as a carbon source. Such metabolic flexibility could provide a local fitness advantage during infection, particularly in SA-rich microenvironments generated at infection sites.

Moreover, it is feasible that local catechol production via SA degradation may contribute to fungal virulence. It has been reported that catechol can autoxidize to a quinone, thereby increasing reactive oxygen species (ROS) production by generating superoxide radicals (Pinnataip & Lee, 2021), which are typically produced by plants during the defense’s oxidative burst (Wojtaszek, 1997). While a common strategy against diverse pathogens, some organisms can exploit it for their own benefit (Di Pietro & Talbot, 2017). This is the case of *B. cinerea,* as ROS not only affect the fungus but also damage host tissues, facilitating colonization and lesion development (Rossi et al., 2017; Siegmund & Viefhues, 2016). We therefore propose that ppNahG-mediated SA catabolism may, at least in part, alter the redox environment, thereby promoting ROS-associated cell death and increasing virulence. Direct assessment of catechol production and ROS dynamics will be required to test if this is the case. Alternatively, a recent report indicates that SA accumulation can influence the physicochemical properties of plant tissues: SA can perturb apoplastic pH homeostasis, leading to alkalinization by inhibiting plasma membrane H⁺-ATPase activity. Notably, this is independent of the NPR1-mediated signaling required for *PR1* induction (Müller et al., 2025). Apoplastic pH is a critical environmental variable for pathogen success, and *B. cinerea* actively promotes host tissue acidification, enabling plant degradation and virulence (Billon-Grand et al., 2012; Müller et al., 2018). In this context, degradation of SA by a SH enzymatic activity has the potential to ameliorate SA-associated alkalinization, thereby restoring a permissive acidic environment that favors fungal colonization.

Despite our findings and proposed hypotheses, the strategies employed by necrotrophic phytopathogens, particularly *B. cinerea*, to shape SA dynamics during plant infection have been largely overlooked. Importantly, the impact of the phytohormone on disease development depends strongly on the pathogen’s lifestyle: while SA generally enhances resistance to biotrophic and hemibiotrophic pathogens, it can be neutral or detrimental when plants encounter necrotrophs, which often exploit host cell death (Zhang et al., 2025). Consistent with this, several phytopathogens, regardless of their trophic strategies, encode functional SHs that modulate SA availability. For example, the necrotroph *Fusarium oxysporum* is directly inhibited by SA, which reduces hyphal growth, sporulation, and virulence, and therefore, deletion of FoSAH1 increases fungal sensitivity to SA, attenuating virulence (Li et al., 2022). Likewise, the hemibiotrophic *Fusarium graminearum* strongly induces the expression of the SH FgShy1 during infection, although its deletion does not affect disease severity in wheat (Hao et al., 2019; Rocheleau et al., 2019). In *S. sclerotiorum*, a closely related necrotrophic fungus to *B. cinerea*, the *ssshy1* gene, which encodes a functional SH, is upregulated during infection, contributing to reduced SA levels and promoting virulence (He et al., 2023). In the maize pathogen *Cochliobolus heterostrophus*, catabolism of exogenous SA is not mediated by an SH but instead occurs through a 5-salicylate hydroxylase that converts SA into gentisic acid. Mutants lacking the enzyme display reduced virulence, elevated SA levels in leaves, and induced expression of SA-responsive genes, indicating that fungal SA metabolism promotes infection (Xu et al., 2025). Similar capabilities have been reported in biotrophic and hemibiotrophic bacteria, including *Candidatus liberibacter* and *Ralstonia solanacearum*, suggesting a convergent pathogenic strategy (Li et al., 2017; Lowe-Power et al., 2016).

The diversity of outcomes mentioned above underscores the need to understand SA metabolism in *B. cinerea*, given its necrotrophic lifestyle, and the paradoxical evidence around SA-related responses. *B. cinerea’s* broad host range may underlie the lack of consensus, and therefore, regulation of SA content during plant infection might be tightly controlled by this pathogen, depending on host, tissue, and temporal variables.

Altogether, the presence of four SH-like enzymes in *B. cinerea*, along with functional precedents from other fungi, suggests that this pathogen may possess an inherent capacity to metabolize SA. Their respective transcript levels appear upregulated in *B. cinerea* during the interactions with multiple hosts, raising the possibility that endogenous SA degradation could contribute to virulence. Interestingly, expression profiles differ across hosts, supporting our previous assessment that SA catabolism by *B. cinerea* may depend on the plant being infected. Moreover, variations in expression levels across time points suggest dynamic modulation of SA levels during the colonization process.

In summary, our results demonstrate that granting *B. cinerea* SH activity via a gain-of-function strategy is sufficient to enhance fungal growth and virulence in hosts with a wild-type SA biosynthetic capacity, independently of major changes in canonical SA responses. By opting for a pathogen-centered approach and conferring SH activity to it, we eliminated potential systemic effects of SA depletion on the whole plant, as previous experiments have done. Moreover, our data supports the idea that *B. cinerea* is capable of modulating SA dynamics, revealing an additional layer by which this hormone can shape disease outcomes. The identification of multiple SH–like genes in *B. cinerea* and their expression patterns during infection suggests that finetuned SA accumulation may constitute an intrinsic virulence strategy in this necrotrophic fungus. Together, these findings support a model in which SA acts not only as a defense signal but also as a local constraint on pathogen colonization, the removal of which favors infection. This work establishes the foundation for future studies addressing the enzymatic and metabolic dimensions of SA degradation in plant–necrotroph interactions.

## MATERIALS AND METHODS

### *Botrytis cinerea* strain and culture conditions

Strain B05.10 of *B. cinerea*, originally obtained from *Vitis vinifera* (Germany), was used as the recipient for genetic modifications (Amselem et al., 2011). For cultivation, regular Petri dishes containing Gamborg B5 (Duchefa Biochemie) supplemented with glucose (20 g/L) were used. The fungus was routinely grown in Percival incubators at 20°C in a 12h:12h light:dark photoperiod. For *in vitro* growth assays, solid Gamborg B5 medium supplemented with sucrose (20 g/L), salicylic acid (1.0 mM), or catechol (1.0 mM) was plated on 15 cm diameter Petri dishes. Plates were inoculated with an agar plug, 5 mm in diameter, containing vegetative non-differentiated mycelium of the B05.10 strain or the herein generated gain-of-function mutants (see below). All plates were incubated at 20°C in a Percival incubator, as mentioned. The area of growth of each fungal colony was measured after 3, 5, and 7 days of cultivation.

### Genetic construct design and *B. cinerea* transformation

The genetic construct for *ppNahG* expression in *B. cinerea* **(Figure S1)** was generated in the linearized pRS426 plasmid containing the URA3 selectable marker using *Saccharomyces cerevisiae* strain BY4741. Yeast recombinational cloning was conducted as described (Oldenburg, 1997) using oligonucleotides listed in **Table S1**. Phusion high-fidelity DNA polymerase was used to amplify the genetic construct. Protoplast-mediated transformation of *B. cinerea* was carried out as described (Schumacher, 2012). Transformants were selected on Gamborg B5-2% glucose containing 50 μg/mL hygromycin B (Invitrogen). Homokaryotic derivatives were obtained through repeated single-spore isolations on Gamborg B5–2% glucose medium with 70 g/ml hygromycin B, followed by transferring individual colonies to fresh Petri dishes. Genomic DNA was extracted from each colony using the Wizard Genomic DNA Purification Kit (Promega), and PCR was performed to confirm the correct insertion of the genetic construct. The PCR-verified mutants were analyzed by qPCR to determine the number of *hph* cassette integrations in the genome. For this purpose, the ratio (in copy numbers) of the *hph* resistance cassette and the single-copy gene *bcfrq1* was determined as described (Vasquez-Montaño et al., 2020). The resulting *hph*/*bcfrq1* ratios were 1.22 and 1.20 for *ppNahG*::OE-1 and *ppNahG*::OE-2, respectively, confirming a single integration event.

### Plant material and virulence assays

*Phaseolus vulgaris* (cv. Apolo, from Instituto de Investigaciones Agropecuarias, INIA-Chile) was cultivated in growth tents for 1 week at 22°C under a 12h:12h light:dark photoperiod before virulence assays were conducted. *Arabidopsis thaliana* Columbia-0 (Col-0), *sid2-2* (Wildermuth et al., 2001), and pPR1:YFP transgenic *A. thaliana* plants (Poncini et al., 2017) were used. *A. thaliana* seeds were soaked for two days at 4°C in darkness. Stratified seeds were potted in a mixture of vermiculite:peat (1:1) and grown for 4 weeks at 22°C under the same photoperiod before use in virulence assays.

For virulence assays, *B. cinerea* B05.10 and the obtained mutants were first grown on Potato Dextrose Agar (PDA) plates supplemented with 100 g/L of blended *P. vulgaris* leaves. These plates were incubated for 1 week as mentioned, and thereafter, used to collect conidia. Asexual spores were collected on liquid Gamborg B5 supplemented with 20 g/L of glucose. The solution was diluted to 2 × 10^5^ spores mL^-1^ containing 10 mM phosphate buffer, pH 6.4. Conidial suspensions (7.0 μL) were used to inoculate leaves of *P. vulgaris* and *A. thaliana*. Lesion size on leaves was determined using ImageJ software (Schneider et al., 2012).

### RNA extraction and RT-qPCR analysis

Infected *A. thaliana* leaves were collected 24 hpi using a 5mm hollow punch tool and immediately frozen in liquid nitrogen. Samples were then lyophilized overnight at -40°C. Leaf tissue was ground into powder, resuspended in TRIzol reagent (Invitrogen), and incubated at 65°C for 5 min. Subsequent steps were performed according to the TRIzol reagent instructions. Before cDNA synthesis, residual DNA degradation was performed using RQ1 DNase (Promega). cDNA synthesis was conducted using 500 ng of total RNA employing the MMLV reverse transcriptase (Promega). The resulting samples were used for quantitative polymerase chain reaction (qPCR) using the SSOAdvanced Universal SYBR Green Supermix on a CFX96 Real-Time System (Bio-Rad). Oligonucleotides are listed in **Table S1**.

### Microscopy

*A. thaliana* pPR1::YFP reporter line, in which the yellow fluorescent protein (YFP) is under the control of the PR1 promoter, was used as a marker of salicylic acid (SA) pathway activation, allowing visualization of SA-mediated signaling in response to *B. cinerea* infection. Virulence assays were performed as described above, and lesion images were analyzed at 24 hpi using an epifluorescence Olympus-IX81 microscope. The Q-capture Pro7 software was used to acquire the images.

### Bioinformatic analysis

To identify candidate salicylate hydroxylases in the *B. cinerea* genome database (DB), we used the *P. putida* enzyme NahG amino acid sequence as a query for a BLASTp. Four proteins (Bcin06g05670, Bcin05gh08190, Bcin10g05120, and Bcin02g00013) with identity over 25% were identified and aligned using Clustal Omega via the EMBL-EBI job dispatcher tool (Madeira et al., 2024; Sievers & Higgins, 2021). A phylogenetic tree and bootstrap analysis (10,000 samples) were constructed using IQ-TREE software version 2.4.0 (Minh et al., 2020) and visualized in FigTree version 1.4.4 (2025). Identification of functional domains was performed by searching the InterPro DB using the amino acid sequences of each candidate enzyme from *B. cinerea* and ppNahG (Quevillon et al., 2005). To visualize the enzymatic active site among the identified proteins, predicted structures computed in AlphaFold were used (Jumper et al., 2021). For this purpose, 3D protein models were visualized employing ChimeraX 1.9 software (Meng et al., 2023).

Gene expression levels of candidate SH-encoding genes were calculated from publicly available transcriptomics data. For this purpose, dual RNA-Seq data from *B. cinerea* infecting *A. thaliana*, *S. lycopersicum,* and *Lactuca sativa* (NCBI SRA accessions PRJNA1158549, PRJNA628162, and PRJNA1013336, respectively) (Pink et al., 2023; Srivastava et al., 2020; Wei et al., 2024) were downloaded and used for Z-score visualization of expression levels. Succinctly, low-quality reads and adapter sequences were filtered from FASTQ files using BBDuk (https://sourceforge.net/projects/bbmap/) according to the developers’ instructions. Gene expression in the three datasets was determined by pseudoalignment of reads to the *B. cinerea* transcripts using Kallisto software, with default settings (Bray et al., 2016). Details of the bioinformatic pipeline have been described elsewhere (Olivares-Yañez et al., 2021). Finally, the obtained gene expression matrices were used as input for ClusterGVis analysis for heatmap generation and visualization (Zhang et al., 2025).

## ACKNOWLEDGMENTS

We acknowledge Danae Ramirez and Sandy Rojas for technical assistance with the cultivation of *Phaseolus vulgaris* and *Arabidopsis thaliana*. This research was funded by the iBio Institute, Iniciativa Cientifica Milenio-MINECON, and ANID/FONDECYT 1240742 to P.C. and ANID/FONDECYT 1240785 to A.H.-V.

---

**Figure S1.**
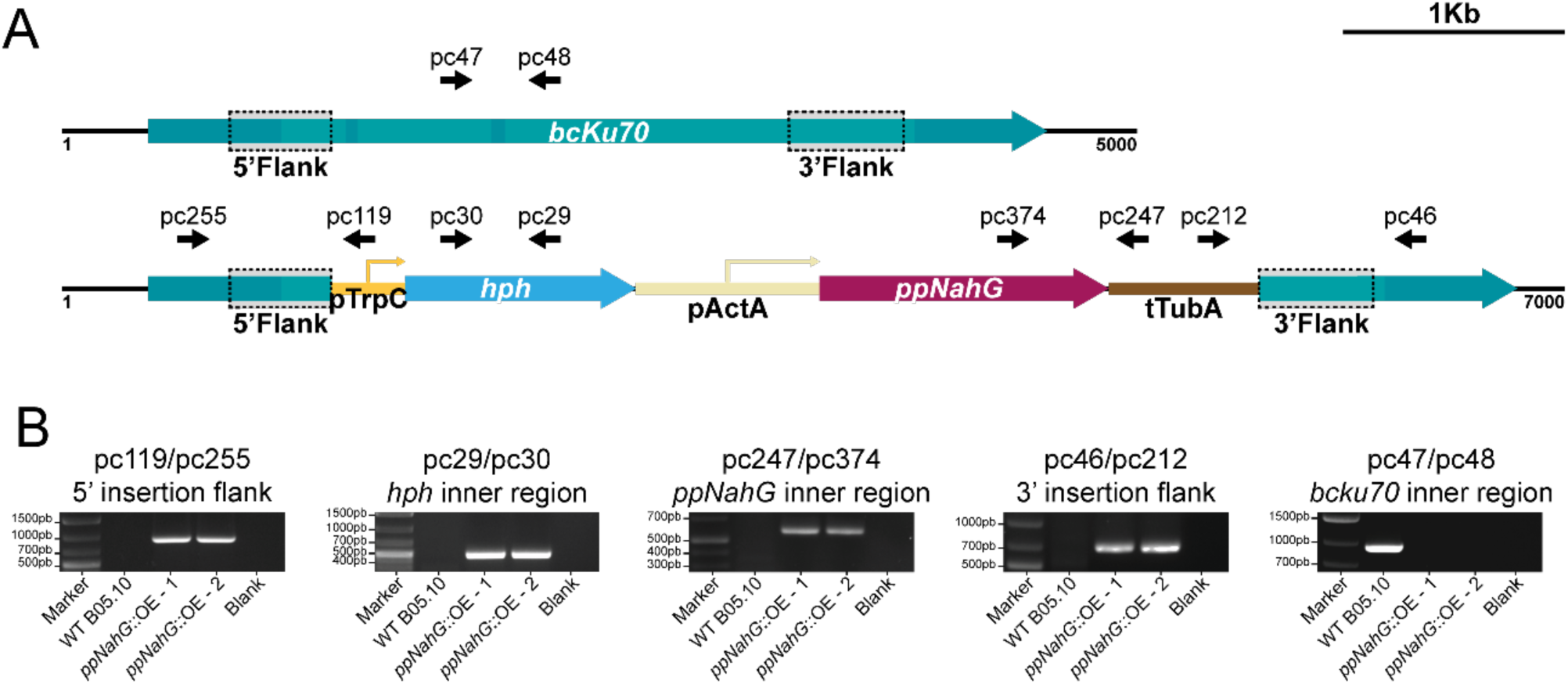
Transformation strategy for *ppNahG* expression in *Botrytis cinerea*. **(A)** Scheme depicting the homologous integration of the *ppNahG* expression construct (bottom) targeted to the *bcku70* locus (Bcin06g03990) of *B. cinerea* (top). Expression of *ppNahG* was driven by the *bcactA* promoter, and transformants were selected for hygromycin resistance (*hph*). Arrows denote transcriptional orientation. Primers used for genotyping **(Table S1)** are indicated with black arrows indicating orientation. The 5’ and 3’ recombinational flanks allowed homologous recombination in the *B. cinerea* genome. **(B)** Genotyping of transformant clones was performed using genomic DNA and compared with the wild-type B05.10 strain by PCR. The PCR amplicons were visualized by agarose (1%) gel electrophoresis.

**Figure S2.**
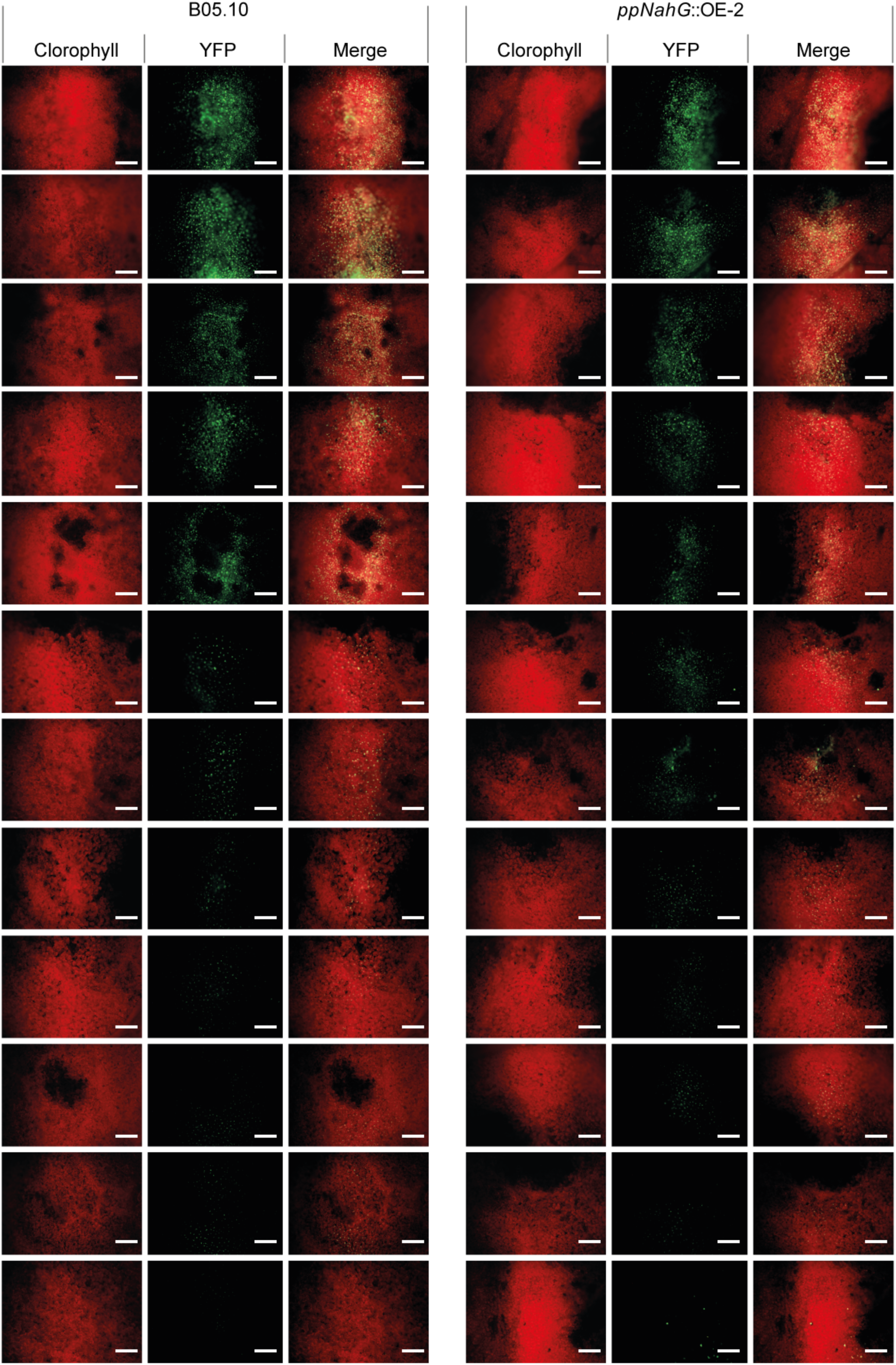
Expression of *ppNahG* in *Botrytis cinerea* does not affect *PR1* promoter activity during infection. Epifluorescence microscopy of *A. thaliana* pPR1::YFP leaves at 24 hours post infection (hpi) with *B. cinerea* B05.10 and *ppNahG::OE-2*. YFP fluorescence is shown in green, and chlorophyll autofluorescence in red. Scale bar = 50µm. Images of proximal and distal regions relative to the lesion site are shown. Across the 12 rows, an equal number of infections (leaves) are observed for each genotype under analysis. The obtained images were arranged from highest to lowest YFP signal (top to bottom, respectively).

**Figure S3.**
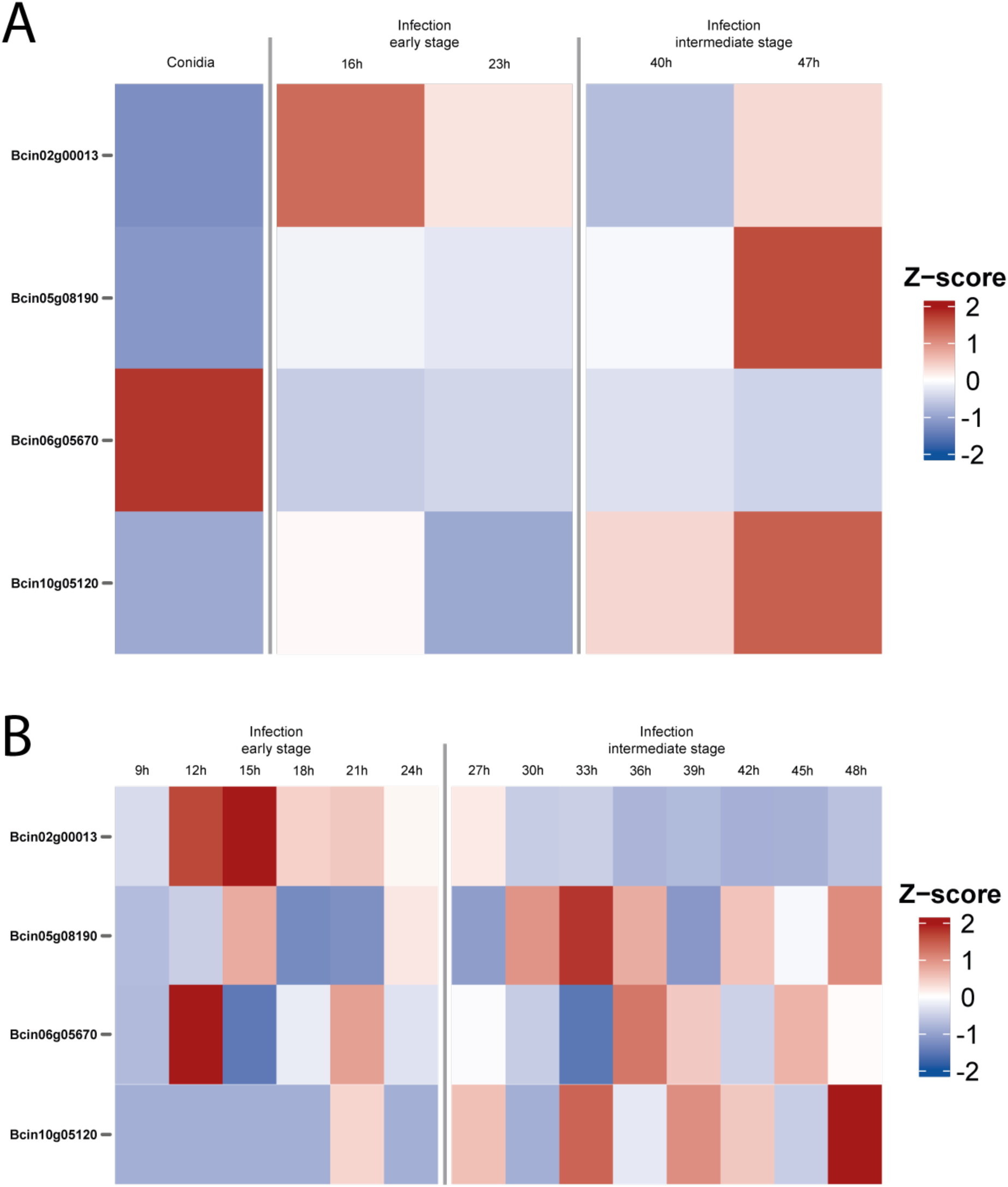
Expression profiles of putative SH enzyme-encoding genes identified in the *Botrytis cinerea* genome. Heatmaps illustrating the relative mRNA levels of putative SH enzyme-encoding genes identified in *B. cinerea* during *S. lycopersicum* **(A)** (Srivastava et al., 2020) and *Lactuca sativa* **(B)** (Pink et al., 2023) virulence assays, at the indicated time points (h). Expression profiles were obtained and calculated from NCBI’s SRA accession numbers PRJNA628162 and PRJNA1013336, respectively.

**Table S1.**
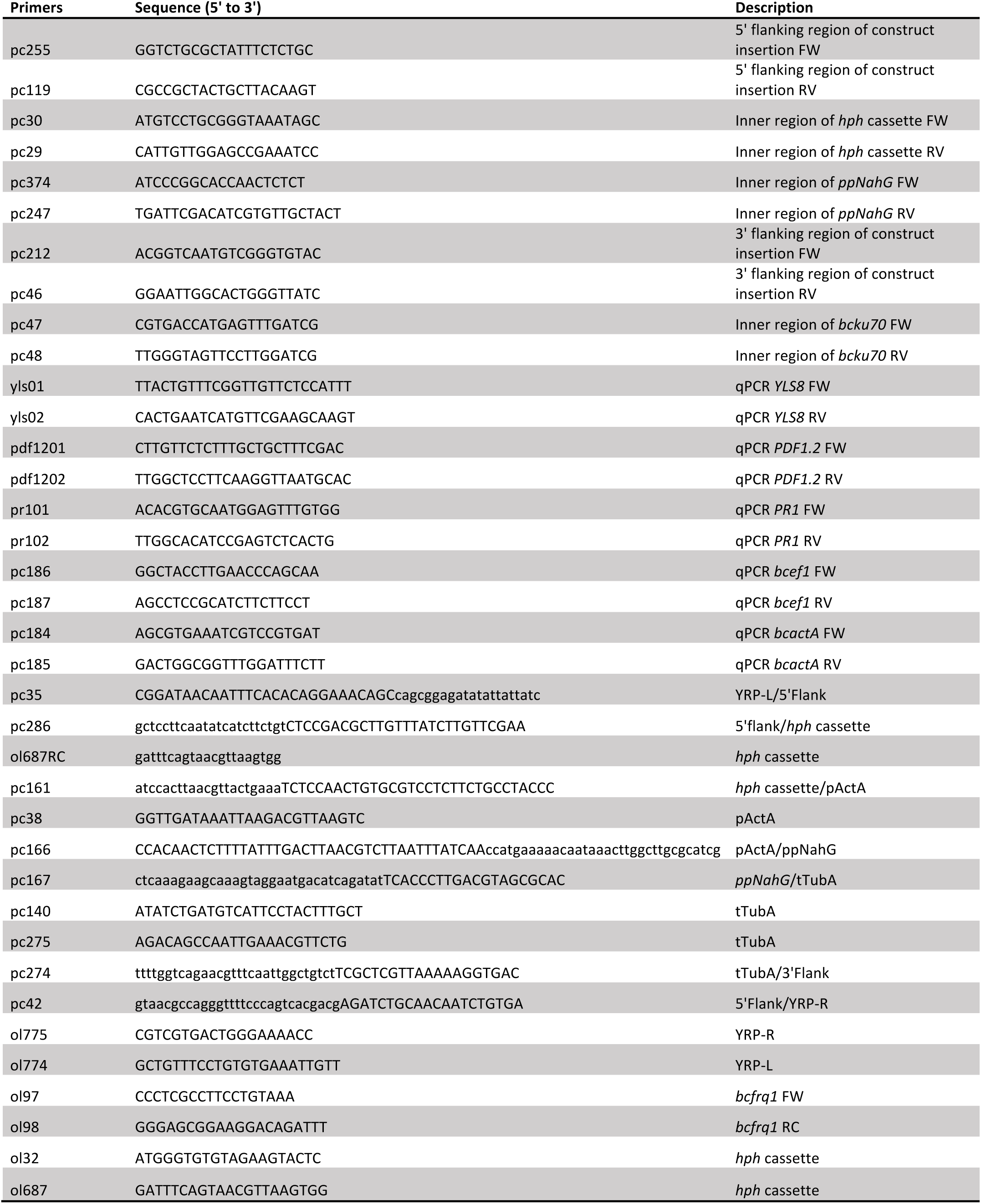
Primers used for molecular analyses in this study. Oligonucleotides used to verify the correct integration of genetic constructs in *B. cinerea* and for RT-qPCR assays are listed. Primers employed for genetic construct assembly are also included.

